# Dynamic multi-omics analysis of Adenovirus-induced rAAV-producing stable cell lines reveals the interplay between adenoviral infection and host cell metabolism

**DOI:** 10.64898/2026.11.12.766245

**Authors:** Saratram Gopalakrishnan, Eunil Im, Leo Dworkin, Madhuresh Sumit, Yingchao Nie, Amit Mathur, Karen A. Vincent, Bruno Figueroa, Nathan E. Lewis

**Affiliations:** Department of Pediatrics, University of California San Diego; Department of Bioengineering, University of California San Diego; Genomic Medicine Unit CMC, Global CMC Development, Sanofi R&D, 225 2nd Ave Waltham, MA 02451

## Abstract

Recombinant adeno-associated viruses (rAAV) are invaluable delivery vehicles for gene therapy. However, to make rAAV-based therapies more accessible, there is a need to increase rAAV titers. Here we analyze the phenotypic variation among 30 Adenovirus-induced stably-transfected human producer cell clones to identify metabolic signatures associated with high vs low rAAV production. Exometabolomic characterization revealed decreased amino acid consumption in higher producers. Analysis of RNA-Seq data in the context of a genome-scale metabolic model identified upregulated carnosine biosynthesis-associated pathways during the first 6 hours post infection with adenovirus, relative to uninfected growth. Meanwhile, late infection timepoints exhibited increased amino acid release and *de novo* biosynthesis of serine, glycine, and arginine. Moreover, high producers decreased expression of genes linked with cholesterol biosynthesis and increased HMG-CoA degradation via HMGCL. Furthermore, sphingosine and ceramide biosynthesis genes were altered in high-producers in directions previously reported to resist cell death and suppress immune signaling. Overall, these findings elucidate the metabolic modulation upon adenoviral infection in a helper virus-induced AAV production process using stably transfected human cell lines.

## Introduction

Recombinant adeno-associated viruses (rAAVs) are a popular vector for gene therapy delivery due to their low pathogenicity, cell cycle independent infection, and capacity for sustained transgene expression. A highly productive biomanufacturing infrastructure is necessary to meet the growing demand for rAAV-based gene therapies. However, the industry-standard production processes face considerable scalability and cost-effectiveness hurdles stemming from low vector yields, reliance on transient transfection of multiple plasmids, and product heterogeneity arising from inefficient capsid packaging and host cell protein contamination. These factors, in conjunction with strict regulatory compliance and complex multi-step workflows further exacerbate accessibility of these therapeutic modalities. Although some of these challenges may be mitigated using stably-transfected cell lines, rAAV production must still be induced (e.g., using a helper virus such as adenovirus 5 (Ad5)), leading to the potential for further product contamination and competition for intracellular resources^1,2^. Therefore, there is a need to mechanistically characterize the cellular machinery governing rAAV production, including the dynamics, metabolic shifts, and resource allocations that govern rAAV genome and capsid synthesis, and the metabolic bottlenecks that must be alleviated to improve productivity and product quality^3–5^.

Multi-omics analyses are transforming biomanufacturing by providing mechanism-based insights into process bottlenecks. These workflows synergistically combine multiple types of -omics data, (e.g., genomics, transcriptomics, proteomics, metabolomics, and fluxomics) and algorithms to elucidate the interplay between different biological networks, thereby revealing actionable engineering strategies aimed at improving productivity and quality^6,7^. Multi-omics analyses have extensively been applied to characterize and engineer other cell factories, such as CHO-based biologics production. Analysis of time-course transcriptomics, metabolomics, and fluxomics data in CHO cells has identified amino acid utilization bottlenecks, particularly arginine, proline, asparagine, and glutamine, that offer actionable targets to boost titers^8^. By forecasting amino acid usage and potential depletion, multi-omics analyses enable predictive control of CHO-based processes^9^. For quality assessment and control, multi-omics has helped elucidate the effects of culture pH^10^ conditions and intracellular metabolic fluxes^11^ on protein glycosylation. More recently, integrated omics analyses have been deployed to study the effects of ER stress activation on the proper assembly of multi-specific antibodies^12^.

For rAAV biomanufacturing, multi-omics analysis has been applied primarily to HEK293 triple transfection processes. A time-course analysis using a genome-scale model revealed HIF-1α as a potential inactivation target that suppressed cell growth and increased critical capsid synthesis but decreased viral genome synthesis through the inhibition of the oxidative pentose phosphate pathway^13^. Another study explored the effects of small molecule additives to the culture medium and found up to 50% increase in rAAV titers using inhibitors of NF-κB, PKC, and TBK1/IKKe^14^. To decrease the reliance on transient transfection, an inducible stably transfected line of HEK293 was constructed and reported a drop in the rAAV capsid titer by an order of magnitude. This loss was attributed to several factors including Rep/Cap expression, rAAV replication kinetics, capsid production limitations, inflammation, immune responses, and MAPK signaling pathways^15^. On the other hand, helper virus-dependent producer cell lines have been constructed and are being explored for commercial applications^1^. These systems mimic the natural replication cycle of AAV by inducing rAAV production via infection with a wild-type adenovirus helper (Ad5), although, resource allocation between different products (viz. Ad5, rAAV, host cell biomass, and metabolic byproducts) remains poorly understood and yield limitations may arise from competition by Ad5 synthesis mechanisms. This motivates a thorough multi-omics analysis to elucidate the biosynthesis pathways, metabolic shifts, and resource allocations between high and low producer cell line-based production systems.

Here we explore the dynamics and variances in metabolism and resource allocation in 30 AAV producer cell line (PCL) clones engineered to produce the same AAV vector (identical capsid and transgene) upon infection with Ad5. A partial least-squares regression analysis of overall metabolite consumption and product formation revealed a negative correlation between rAAV titer and essential amino acid consumption. A lower amino acid uptake was also associated with decreased Ad5 titers. A closer inspection of uptake and secretion rates during the growth phase, early infection phase (0hpi – 6hpi), mid-infection phase (6hpi – 24hpi), and late infection phase (24hpi – 48hpi) revealed an increase in growth rate immediately upon infection followed by a sharp reduction in the uptake of essential amino acids after 6hpi. Analyzing the transcriptomic profiles between high and low producers post infection revealed an up-regulation of pathways associated with amino acid storage during early infection and increased *de novo* amino acid synthesis during the late infection in high producing clones. Although the transcript abundances for the primary amino acid transporter LAT1 did not significantly change, metabolic changes were seen in cholesterol biosynthesis, lipid rheostat, and genes associated with Golgi transport regulation. Overall, these findings reveal the cell-wide impacts of adenoviral infection and highlight potential cell line engineering targets that can modulate amino acid metabolism and intracellular signaling to improve the productivity of rAAV, which can lower manufacturing costs and increase accessibility to gene therapy treatments.

## Results

### Clonal rAAV productivity is inversely correlated with essential amino acid consumption

Thirty clones derived from three different parental PCL pools (MW1, MW2, and MW3) show a 10-fold range in rAAV titer and a 2.5-fold range in Ad5 titer post-infection with Ad5 (Figure 1B and C). The top-producing clones were derived from MW1 whereas MW3 had the poorest rAAV producers. Clones from MW1 showed significantly higher rAAV titers (p < 0.025 rAAV; Mann-Whitney U-test) and lower Ad5 titers compared to the other PCLs and the parental control (p < 0.05 Ad5; Mann-Whitney U-test). However, clones derived from MW2 and MW3 showed minimal differences (p > 0.15; Mann-Whitney U-test).

**Figure 1:**
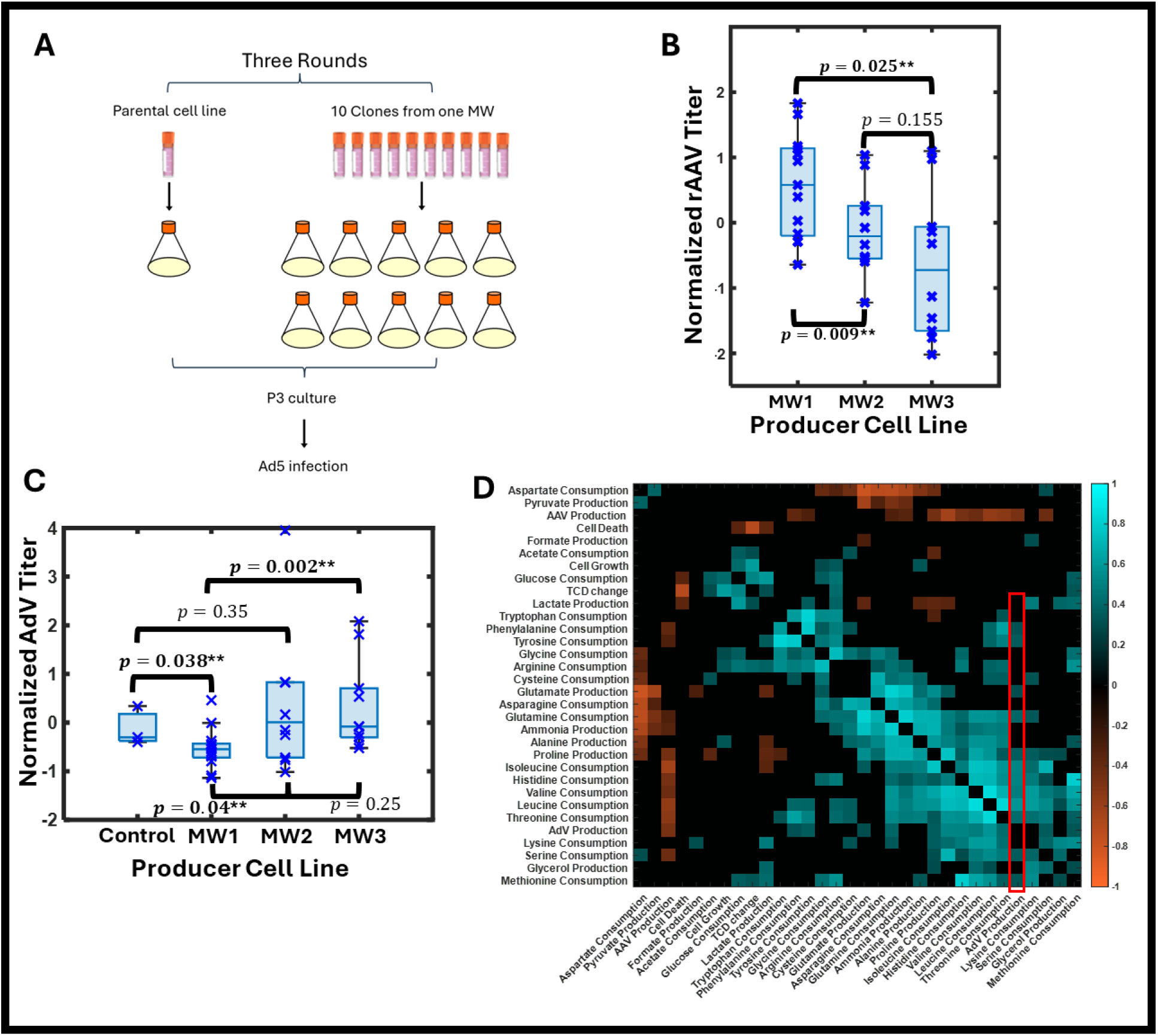
Overview of the rAAV production process. (A) The experimental setup showing the steps in rAAV manufacturing. Stably transfected parental cell lines are transferred to a production medium and infected with Human Adenovirus 5 (AdV) to induce the simultaneous production of AdV and rAAV. (B) Box and whisker plot showing the range of rAAV production among the three parental pools (MWs) each with 10 clones considered in this study. Normalized rAAV titers represent the z-score. Significant p-values for the one-tailed Mann-Whitney test are shown with an asterisk. (C) Box and whisker plot showing the range of AdV titers among the parental control and the three MWs. Normalized AdV titers represent the z-score. Significant p-values are indicated with an asterisk. (D) Heat map of Spearman correlation coefficients shown for pairs of measured quantities with p-value less than 0.05. Rows and columns are ordered based on Euclidean distance-based clustering. The highlighted red box shows the amino acids positively correlated with AdV titers.

We next investigated how various factors co-vary and correlate with virus titers. The Spearman correlation coefficient (SCC) of the metabolite consumption/production over the course of viral vector production and the final Ad5 and rAAV titers (Figure 1D) showed that a higher rAAV titer was associated with reduced essential amino acid consumption, particularly glutamine, histidine, aromatic amino acids, and branched chain amino acids. This suggests that the best rAAV producers consumed fewer essential amino acids overall. This negative correlation extended to other amino acids as well but were not statistically significant (p >= 0.05) (Supplementary Figure S1). Ad5 production, however, positively correlated with the consumption of essential amino acids, suggesting that decreased Ad5 titers in high-producing clones is mediated by decreased amino acid channeling towards Ad5 biosynthesis.

To elucidate the trade-offs between the key products (cell biomass, lactate, alanine, proline, glycine, Ad5, and rAAV), we conducted partial least squares (PLS) regression using the normalized metabolite consumption, cell density, and the Ad5 titer as the predictor and the normalized rAAV titer as the response. The number of PLS components was set to seven as this was the minimum number required to explain 95% of the observed variance in rAAV titer across the 30 clones (Supplementary Figure S2A). The PLS model using seven components showed good agreement between the predicted and observed rAAV titers (R^2^ = 0.97, Supplementary Figure S2B). Because the consumption/production of several metabolites, notably essential amino acids, tends to be highly correlated in cell culture processes, Variable Importance in Projection (VIP) scores were computed to find predictors underlying the observed variances^16^ (Figure 2A). The 14 predictors with a VIP score greater than 1 included seven essential amino acids, three non-essential amino acids, ammonia, two central metabolites, and Ad5. Despite fluctuations over the course of the production process, changes in total cell density did not present itself as a significant predictor. For a deeper understanding of the directionality of influence of each significant predictor, we visualized the PLS predictor scores projected onto the first two PLS components. Increases along each individual PLS component were associated with an increase in rAAV titer. PLS component 1 explained 26.8% of the total variance in the predictors and primarily captured the differences between MW1 and MW3 and reflected changes in histidine, valine, leucine, and threonine consumption. Meanwhile, PLS component 2 captured the differences between MW1 and MW2, and captured differences in glutamine and serine consumption. Given the positive correlation between Ad5 titers and amino acid consumption, Ad5 production negatively influenced both PLS components. Overall, these findings indicate a suppression of amino acid metabolism in conjunction with Ad5 synthesis as a key metabolic feature of high-producing clones.

**Figure 2:**
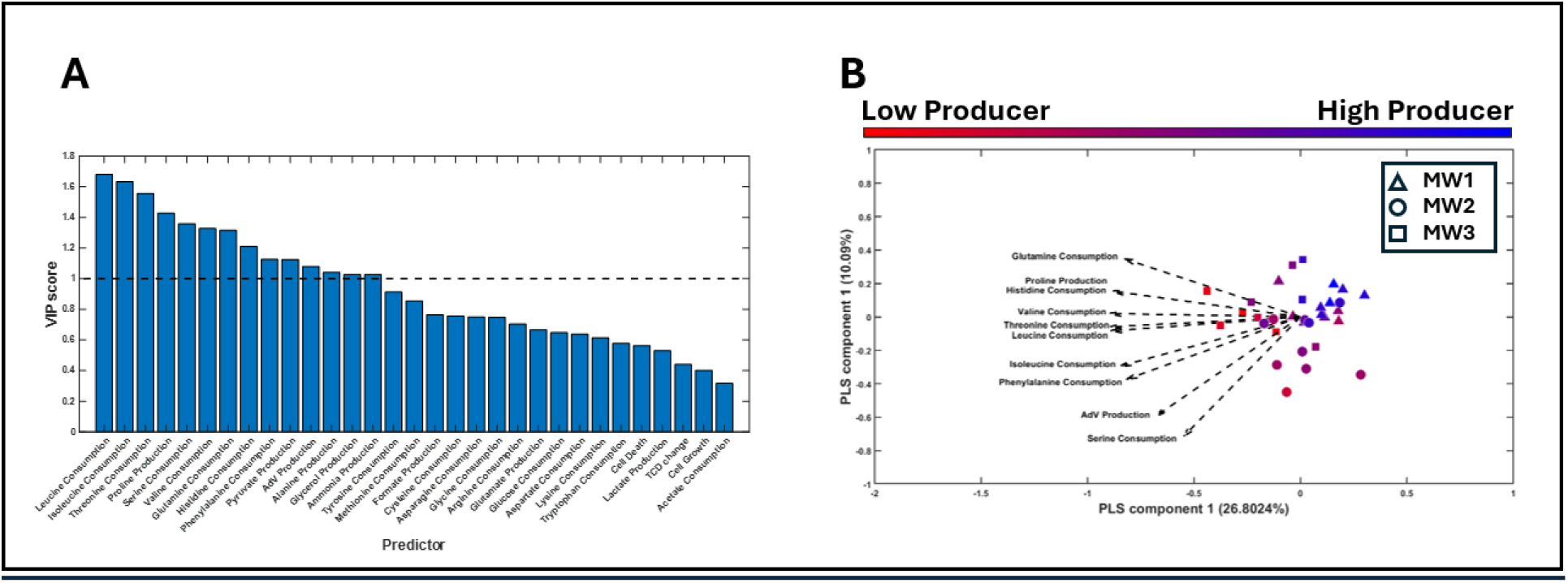
Partial Least Squares (PLS) regression analysis showing the factors that are correlated with increased rAAV production among the 30 clones of the three PCLs. (A) Bar graph showing the Variable Importance in Projection (VIP) scores for the 31 predictors used for PLS regression. Predictors with a VIP score greater than 1 are considered important. (B) Scatter plot showing the predictor data matrix projected on to the first two PLS components. Different markers represent different PCLs and the color represents the relative rAAV titer from least (red) to most (blue). The embedded biplot show the relative weight of the top 10 factors driving the observed variances among the 30 clones.

### Clonal variation in metabolism is prominent in the early-infection phase

To understand the metabolic basis for the observed variations in amino acid utilization, we performed a PCA on the normalized computed uptake and secretion rates (see Methods) (Figure 3B). The observations separated into four clusters based on process stage, including growth (pre-infection), early infection (0 – 6 hpi), mid-infection (6 – 24 hpi), and late infection (24 – 48hpi). The second principal component captured the metabolic shifts between growing cells and early infection. This metabolic shift entails a four-fold increase in the median lactate secretion, a three-fold increase in glucose uptake, a two-fold increase in the uptake of most essential amino acids, a reduction in cell division, and a 40% reduction in median asparagine uptake. The second metabolic shift was observed at 24 hours post infection and is represented by a leftward shift along principal component 1. This shift was characterized by a 75% reduction in amino acid uptake/secretion, cell death, and a reduction in glucose uptake. Interestingly, lactate secretion increased further in this phase. Since the late-stage adenoviral genes are typically expressed 8 hours post infection^17^, and rAAV rep/cap amplification begins 12 - 24 hours post infection depending on the producer cell line^18–20^, it can be inferred that most virus particles (both Ad5 and rAAV) are synthesized during the late infection phase. To further investigate whether the amino acid and glucose uptake in the late infection phase was sufficient to meet the metabolic demands for Ad5 and rAAV synthesis, we performed a Flux Balance Analysis using the human metabolic model, Recon2.2^21^ after including macromolecular reactions for Ad5 and rAAV biosynthesis and imposing the computed uptake and secretion rates as boundary flux constraints. During this phase, extracellular amino acid uptake accounted for less than 10% of the biosynthetic demand for virus production (Figure 3C). Furthermore, glucose uptake was insufficient to explain the observed lactate flux. This suggests that the cells primarily rely on internal reserves such as proteins and glycogen as a source of precursors to meet the metabolic demands in this phase.

**Figure 3:**
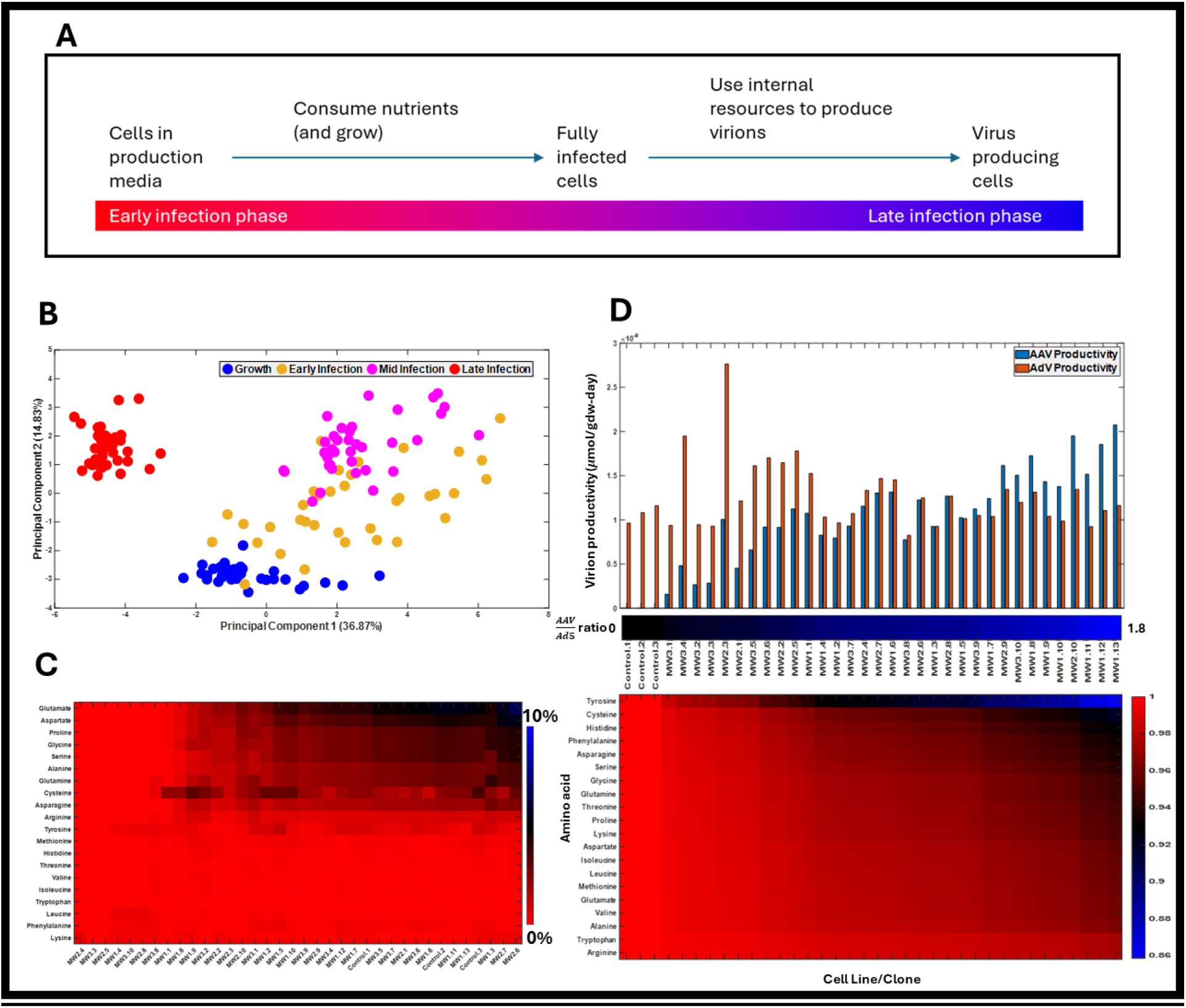
Overview of metabolic shifts and resource allocation in an rAAV manufacturing process. (A) AdV infection induces metabolic shifts that temporally separate uptake and utilization of taken up substrates. (B) PCA scatter plot showing the shifts in metabolite uptake and secretion rates over the course of the bioprocess. Cells in their natural growing state are shown in blue, early infection phase in orange, mid-infection phase in magenta, and virion producing cells in red. (C) Heat map showing nutrient deficit for virion production. A value of 1 (red color) indicates that all nutrients are derived from internal stores. (D) Plot showing the correlation between nutrient distribution towards each virion and rAAV productivity.

A key question that remained to be answered was whether higher rAAV titers seen in some clones was a consequence of increased combined virus production (rAAV + Ad5) or metabolic reprogramming to increase precursor channeling specifically towards rAAV production. Thus, we computed resource allocation towards rAAV and Ad5 among the 30 clones and found that the fraction of amino acids channeled towards rAAV biosynthesis correlated with the specific productivity of rAAV (Figure 3D). However, the productivity of Ad5 among clones did not change significantly, indicating that metabolic reprogramming plays a key role in increasing rAAV titers in high-producing clones.

### High-producing clones increased *de novo* amino acid biosynthesis and ATP production

The observed temporal changes in uptake and secretion rates suggest an underlying transcriptomic and metabolic shift. Time-course RNA-Seq data provided an insight into the transcriptomic shifts underpinning the observed phenotypic changes. Gene expression data was thresholded using the min-max-mean approach and then overlaid on to Recon2.2 using a modified mCADRE method^22,23^ to construct context-specific metabolic networks representing the ensemble of likely active metabolic pathways at each of the six time points evaluated in this study (growth, 0 hours post infection (hpi), 3 hpi, 6 hpi, 24 hpi, and 48hpi; Figure 4A). A comparative analysis of network topologies across each time point revealed 11 pathways that are always retained, 23 pathways that are always inactive, and 18 pathways that showed strong activation or inactivation at one or more time points. A detailed description of perturbed pathway topologies is provided in the Supplementary Results. Notably, taurine metabolism, eicosanoid metabolism, and some nuclear transporters were inactivated at 48 hpi. The biosynthesis of triacylglycerols, sphingolipid metabolism, and degradation of phenylalanine, tyrosine, and histidine were inactivated earlier in the process (0 – 6 hpi). CARNS1 expression, required for synthesizing carnosine from histidine and beta-alanine was higher during the early infection phase, whereas CNDP2, a cytosolic dipeptidase, was more commonly seen in the 48 hpi models. These topological shifts suggest a preservation and storage of consumed amino acids as well as perturbation to lipid metabolism and signaling pathways.

**Figure 4:**
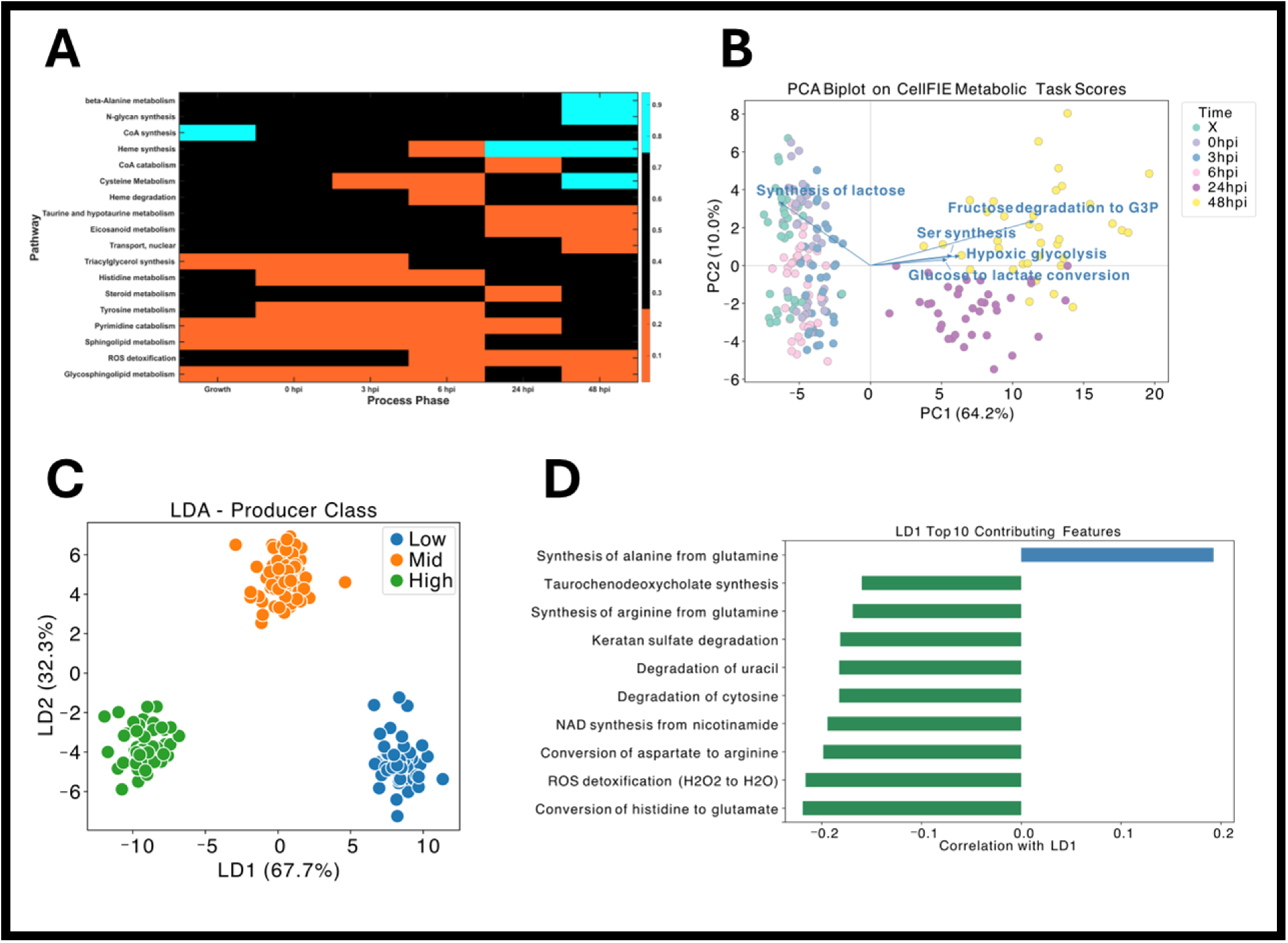
Temporal RNA-Seq profiling of metabolic changes during Adenoviral infection. (A) Topological changes induced by Ad5 gene expression. The heat map indicates pathway retention, which represents the fraction of the pathway that is included in the context specific model. A pathway retention of 25% indicates that 25% of the reactions in the pathway are retained in the context-specific model. (B) PCA of the metabolic task scores computed by CellFIE. (C) LDA performed to identify the tasks that distinguish high and low producers of rAAV. High and low producers include the top and bottom 30 percentile of rAAV producers, respectively. The middle 40 percentile samples represent the mid producers. (D) Correlation coefficients between individual task scores and the first linear discriminant axis (LD1). Tasks with a negative coefficient positively correlate with rAAV titer.

Topological comparison is sensitive to pathways gained or lost between conditions but not to quantitative changes within pathways retained throughout. To elucidate pathway-level differences in conserved topologies, we used the CellFie algorithm^24^. CellFie organizes key reactions within Recon2.2 into 195 curated metabolic tasks that represent biologically observed metabolic functions in human cells. Metabolic tasks spanned key cellular processes that were not significantly topologically altered such as glycolysis, oxidative phosphorylation, amino acid biosynthesis, amino acid degradation, fatty acid biosynthesis and degradation, cholesterol metabolism, and glycan biosynthesis. RNA-Seq data from the growth stage, 0hpi, 3hpi, 6hpi, 24hpi, and 48hpi was overlaid on to genes associated with these metabolic tasks to compute task scores that elucidate aggregated pathway usage.

PCA of CellFie task scores revealed two prominent clusters representing early and late infection stages separated along principal component 1 (PC1), which accounts for 64.2 % of the total variance in the metabolic task score dataset (Figure 4B). The rightward shift indicates increased glycolytic activity and amino acid synthesis. Metabolic tasks associated with glycolysis, such as fructose degradation to glyceraldehyde-3-phosphate (G3P), hypoxic glycolysis, and glucose to lactate conversion exhibited ∼1.2-1.5-fold increase from the early to late infection stage. This increase in gene expression-based task scores aligns with the increased lactate secretion and glucose uptake observed at 6 hpi, supporting a shift toward enhanced glycolytic activity during infection. Tasks associated with amino acid synthesis, including serine synthesis and glycine synthesis, were increased by 1.3-1.5-fold from early to late infection. This increase in *de novo* amino acid synthesis is consistent with suppressed extracellular amino acid uptake during late infection stage, where extracellular uptake contributes less than 10% of the biosynthetic demand for virus production. These findings indicate that viral amino acid requirements are met through *de novo* pathways. The metabolic shift from the growth stage through 6 hpi, as well as the shift from 24-48 hpi were primarily along PC2, which accounts for 10.0 % of the total variance. A downward shift along PC2 indicates a decrease in tryptophan depletion and O-glycan metabolism. (Supplementary Figure S3).

To characterize the metabolic differences between high- and low-producing clones from each parental pool, we performed Linear Discriminant Analysis (LDA) to the metabolic task scores. As LDA identifies linear combinations of features that separate predefined categories, we can examine the metabolic pathways that distinguish high-, mid-, and low-producing clones. Distinct clusters corresponding to each producing class were observed along LD1, indicating a metabolic shift across high-to low-producing clones (Figure 4C). Top metabolic tasks contributing to LD1 were associated with oxidative phosphorylation, *de novo* amino acid synthesis, and pyrimidine degradation (Figure 4D). Increased NAD synthesis and reactive oxygen species (ROS) detoxification were observed in high-producers, suggesting enhanced mitigation of redox generated by glycolysis. “Conversion of histidine to glutamate” and “conversion of aspartate to arginine” are metabolic tasks that support internal redistribution of nitrogen and carbon skeletons and *de novo* amino acid synthesis (Figure 4D). This observation aligns with the reduced extracellular amino acid consumption in high-producing clones, suggesting increased reliance on internal metabolic resource mobilization toward virus production. In addition, high-producing clones exhibited increased nucleotide salvage and increased biosynthesis of taurochenodeoxycholate, a taurine-conjugated primary bile acid.

## Discussion

In this study, we performed a comprehensive multi-omics analysis to quantify the trajectory of cellular metabolism upon Ad5 infection and evaluated the various factors and resource trade-offs affecting rAAV production in 30 AAV producer cell line clones. The total Ad5 production over the 48-hour culture period positively correlated with the uptake of several essential amino acids (i.e., methionine, phenylalanine, tyrosine, histidine, valine, leucine, isoleucine, and glutamine). Among the aromatic amino acids, tryptophan uptake was not correlated with adenovirus production, suggesting that its demand for virus capsid and helper protein synthesis is very low. Phenylalanine and tyrosine were independently correlated with Ad5 production. Although human cells can convert phenylalanine to tyrosine via the Phenylalanine Hydroxylase enzyme encoded by the *pah* gene^25^, this gene is poorly expressed in the parental cells (Supplementary Table ST1), leading to independent uptake and metabolism of phenylalanine and tyrosine during the bioprocess. Among the sulfur-containing amino acids, only methionine correlated with Ad5 production.

Exometabolomic analyses revealed a sharp increase in growth rate within the first six hours following Ad5 infection. This was accompanied by a corresponding increase in the uptake of all essential amino acids. This finding is consistent with the previously characterized roles of Ad5 early genes E1A and E4 in the activation of the cell cycle through pRB sequestration^26,27^ and subsequent activation of central metabolism and amino acid metabolism via MYC^28^. High-producers of rAAV exhibited an enhanced growth rate compared to low-producing clones, implicating the susceptibility of cell cycle activation and resistance to apoptosis as hallmarks of high-producers. The later stages (24 hpi and 48 hpi), however, saw a ∼75% decline in amino acid uptake despite no significant change in the expression of key amino acid transporters (SLC7 and SLC38 classes) and non-depleted amino acids in the media. This suggests that amino acid uptake changes are not regulated at the transcriptomic level, but at the translation, translocation, and proteomic levels. The expression of PI4KB was lower in high-producers. This protein can regulate protein trafficking from the cytoplasm to the plasma membrane through the Golgi^29^. Moreover, PI4KB inhibition can also decrease cholesterol transport through non-vesicular transport mechanisms at the ER-Golgi contact sites^30^. High-producing clones also show a lower expression of HMGCoA-synthase, mevalonate kinase, and other enzymes associated with cholesterol biosynthesis (Figure 5A and B). Moreover, PI4KB and cholesterol can facilitate viral replication and the antiviral response, and are therefore also being investigated as a targets for antiviral therapy^31^. Furthermore, cholesterol can also decrease the V_max_ of amino acid transporters^32^ through direct binding^33^ and/or membrane raft organization^33^. These findings, taken together with a decreased expression of cholesterol biosynthesis genes and decreased overall amino acid uptake by high-producers, suggest a potential mechanistic link between cholesterol metabolism and Golgi transport regulation that functions as a switch to modulate the flow of intracellular amino acids towards Ad5 and rAAV biosynthesis.

**Figure 5:**
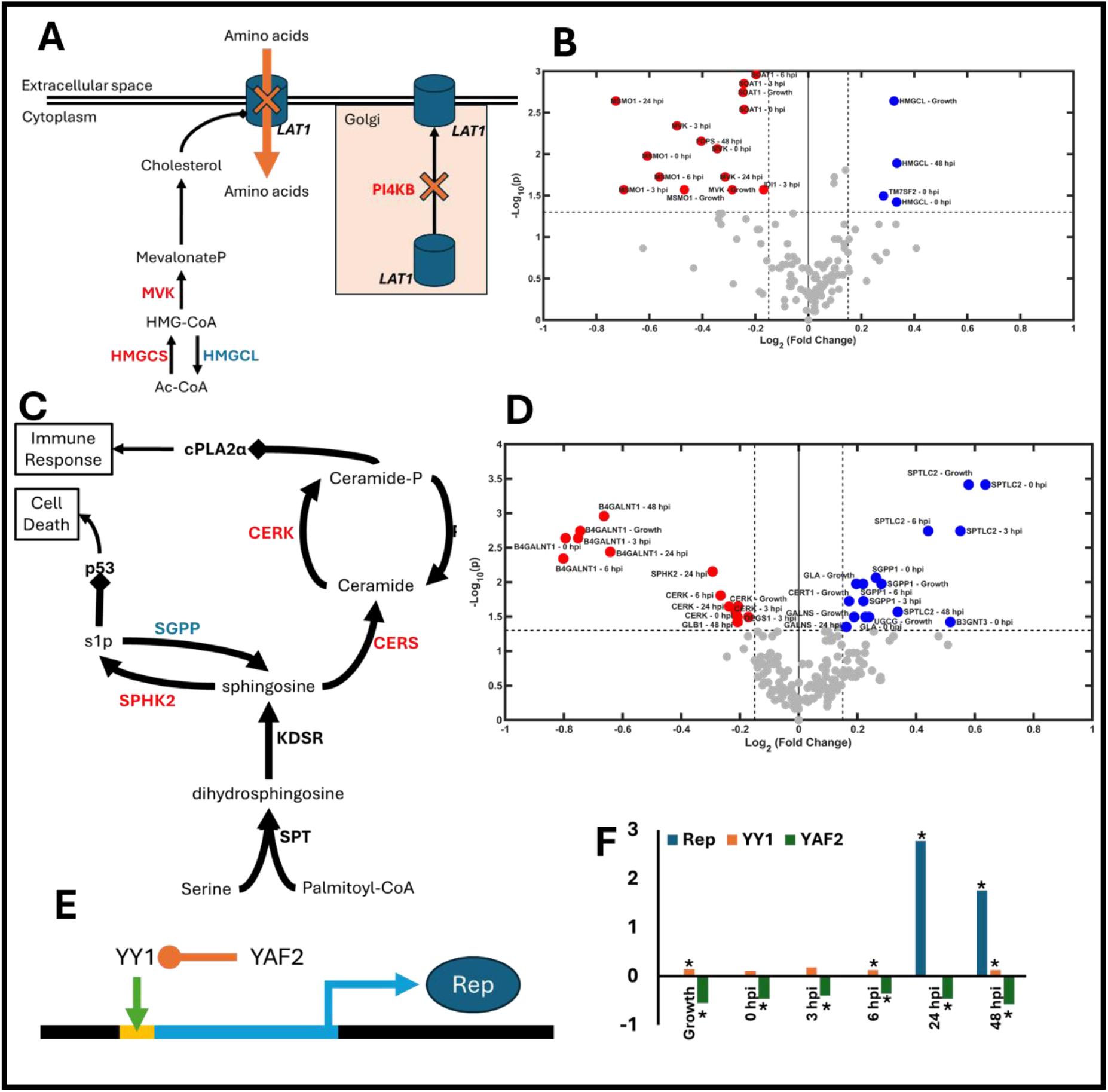
Key genes showing expression differences between high and low producers. (A) Metabolic pathway for cholesterol biosynthesis and its role in stabilizing the amino acid transporter. Genes are indicated in upper case. Down-regulated and up-regulated genes are in red and blue, respectively and the magnitudes and p-value are shown in the volcano plot (B). (C) The sphingolipid pathway with genes shown in upper case. Fold changes and significance are shown in the volcano plot (D). (E) Regulatory effect of YAF2 on the p5 promoter. (F) Fold-change in the expression of YAF2, YY1, and AAV-Rep genes in high producers relative to low-producers. A * indicates a p-value of less than 0.05.

Besides changes to amino acid usage, high-producers showed significant differences in sphingolipid metabolism (Figure 5C and D). The biggest differences occurred around the fates of sphingosine. While SPHK1 was unaffected between high and low producers, the high producers showed decreased SPHK2 expression and increased SGPP1 expression. Phosphorylation of sphingosine by SPHK2 in the ER and nucleus can promote apoptosis^34,35^, suggesting that decreased phosphorylation in conjunction with dephosphorylation of sphingosine-1-phosphate may contribute to improved cell survival during the late stages of adenoviral infection. Ceramide synthesis and phosphorylation via CERS and CERK, respectively, were also decreased in high producers. Ceramide-1-phosphate, the product of CERK, directly activates cPLA2α^36^, initiates selective release of arachidonic acid from membrane lipids^37^, and triggers eicosanoid metabolism resulting in cellular anti-viral immune responses^38,39^. Together, these findings implicate apoptosis resistance and pro-survival factors as hallmarks of high rAAV-producing clones. However, additional studies are necessary to quantify the mechanistic networks involving metabolism and signaling networks that underpin enhanced cell survival and sustained rAAV production.

A hallmark distinguishing feature of high-producers was the increased expression of Rep and Cap genes. Rep/Cap expression requires extrachromosomal amplification^20^ and is influenced by the p5 promoter^40^, CARE element, and the Ad5 L4-22/33K proteins^2^. The p5 promoter is activated by YY1^41^, which is negatively regulated by YAF2^42^. We found that YAF2 was consistently down-regulated at all time points in high-producers compared to low-producers (Figure 5E and F). YAF2 is activated in response to cellular stress and DNA damage and is known to trigger p53-mediated apoptosis pathways^43^. These findings implicate relieved regulation of YY1 and enhanced activation of p5 as an additional mechanism associated with increased rAAV production in high-producing clones.

Overall, the findings here present a dynamic picture of the diverse host cell metabolic responses upon infection by Ad5 and its association with rAAV biosynthesis. While certain parts of metabolism serve a biosynthetic role (e.g., glycolysis for ATP, *de novo* amino acid synthesis, and increased nucleotide biosynthesis), several other pathways are either activated or repressed to either elicit or prevent metabolite-initiated cell signaling pathways. These pathways include amino acid catabolism, cholesterol metabolism, and sphingolipid metabolism, which make up a much larger fraction of metabolic genes than central metabolism. Such metabolic modulation highlights the communicative role played by peripheral metabolism and motivates the development of metabolic pathway-based inference tools to trace metabolite origins. In the context of rAAV biomanufacturing, these indicate potential targets for expression perturbation to boost rAAV titers. Three key components of cellular function not explored in this study are lipidomics, non-metabolic metabolite-protein interactions, and protein-protein interactions. However, this study identified significant changes in lipid metabolism and potential alterations to lipid signaling, which can be further clarified with future lipidomics studies that include intracellular lipid pools and changes to plasma membrane lipid compositions. Secondly, this study revealed the metabolic reprogramming corresponding with alterations in pathways associated with known metabolite signaling functions. Constructing comprehensive databases of context-specific interactions can enable meaningful inferences of pathway and metabolite level perturbations on the outcome of the bioprocess. Finally, metabolism itself is modulated by enzyme levels that are directly impacted by the phases of the cell cycle. To this end, a mechanistic model of relevant protein signaling networks governing cell growth and interaction networks regulating protein transport across the Golgi will be useful to bridge this gap. A quantitative model that accounts for all of these features will be able to predict the dynamics of viral infection and rAAV productivity even under conditions when fluxes are not directly quantifiable using existing methods such as FBA or Monte-Carlo sampling. This will enable digital twin construction for process optimization and cell line engineering to maximize the production of viral vectors for gene therapy.

## Acknowledgements

The authors acknowledge generous funding from Sanofi, which supported this work. The authors would like to thank Sanofi GMU CMC Rebecca Jeppson, Indumathi Subramanian for the experiment set up and sample generation. Thank GMU CMC Bioprocess Analytic group for supporting of AAV and Ad5 titer analysis.

## Methods

### Cell Line Generation and Culture Conditions

AAV producer cell lines (PCLs) were generated as previously described^19^. Parental and AAV producer cell lines were cultured in suspension shake flasks containing ExCell medium (Sigma-Aldrich, Cat#14591C) supplemented with 6 mM L-glutamine (Gibco, Cat#25030081) at 37°C, 100 rpm, 5% CO₂, and 80% humidity. Cells were passaged every 3–4 days for routine maintenance and experimental scale-up.

### Experimental Design and Sample Collection

Thirty clonal producer cell lines from three parental pools (MW1, MW2, and MW3) banked during a single campaign were thawed in three sequential rounds based on parental pool origin. Ten clones from MW3 were thawed in round 1, ten from MW2 in round 2, and ten from MW1 in round 3. A cryovial of parental cell line was thawed and cultured alongside each round as a control.

For AAV production, PCLs were infected with wild-type adenovirus 5 (Ad5). Cells were scaled up in shake flasks, and day 3 post-seeding passage 3 (P3) cultures (N-1 growth phase) were used for infection in 250 mL shake flasks (SF250, N production phase).

### Sampling and Measurements

Cell culture samples were collected at the following timepoints: N-1 days 0, 2, and 3 (pre-infection), and N production at 0, 3, 6, 24, 48, and 72 hours post-infection (hpi). At each timepoint, cell density and viability were measured using ViCELL Blu (Beckman Coulter), and pH was measured using RAPI Point 500 (Siemens).

For spent medium analysis, samples from N-1 day 3 and N production at 0, 3, 6, 24, and 48 hpi were centrifuged at 500×g for 3 min. Supernatants were filtered through 0.2 µm filters and stored at -80°C. Cell pellets from the same timepoints were washed once with cold PBS, snap-frozen, and stored at -80°C for RNA sequencing. Whole cell cultures at 48 and 72 hpi were collected for AAV and Ad5 titer analysis. Samples were stored at -80°C until shipping out in dry ice for analysis.

### Analytical Methods

Spent medium analysis was performed by the Analytical Service Unit (iBET, Portugal). Metabolites (acetate, glucose, formate, lactate, ammonia, and pyruvate) were quantified using a Cedex Bio Analyser (Roche). Amino acid concentrations were measured by HPLC-UV at 260 nm following derivatization using a Waters kit. RNA extraction and sequencing were performed by Genewiz (Azenta).

### RNA extractions and Sample QC

Total RNA was extracted from fresh frozen cell pellets using the Qiagen RNeasy Plus Mini kit following manufacturer instructions (Qiagen, Hilden, Germany). Total RNA was quantified with a Qubit 2.0 Fluorometer (Life Technologies, Carlsbad, CA, USA) and RNA integrity was verified on a 4200 TapeStation (Agilent Technologies, Palo Alto, CA, USA).

### Library Preparation and sequencing

RNA sequencing libraries were prepared with the NEBNext Ultra II RNA Library Prep Kit for Illumina following manufacturer’s instructions (New England Biolabs, Ipswich, MA, USA). Briefly, mRNA was enriched using Oligod(T) beads. Enriched mRNA was fragmented for 15 min at 94 °C. First and second strand cDNA were subsequently synthesized. cDNA fragments were end-repaired and adenylated at 3’ ends. Universal adapters were ligated to cDNA fragments, followed by index addition and library enrichment by PCR with limited cycles. The libraries were validated on an Agilent TapeStation (Agilent Technologies, Palo Alto, CA, USA), and quantified with a Qubit 2.0 Fluorometer (ThermoFisher Scientific, Waltham, MA, USA) and quantitative PCR (KAPA Biosystems, Wilmington, MA, USA). Sequencing libraries were multiplexed and clustered on an Illumina flowcell. After clustering, the flow cell was loaded on a Illumina NovaSeq 6000 and sequenced using a 2×150bp Paired End (PE) configuration and targeting ∼30 million reads per sample. Raw sequence data (.bcl files) were converted into fastq files and de-multiplexed using Illumina bcl2fastq 2.20 software. One mis-match was allowed for index sequence identification.

Gene quantification was done using the GRCh38.p13 release 38 reference genome assembly. Ad5 reference genome and annotation was obtained from NCBI (RefSeq accession AC_000008.1; GenInfo accession 56160529; RefSeq assembly GCA_000857865.1; GenBank assembly GCF_000857865.1; Name ViralProj15107). The AAV gene annotation and reference genome was prepared by Sanofi. Both genome indexing and alignment of trimmed reads were performed by the STAR aligner (v2.7.3a). Prior to alignment, quality control of paired-end fastq reads using FastQC (v0.11.9), Trimmomatic (v0.39). This ensured removal of low-quality and adapter sequences. Following alignment, both raw and transcript-per-million (TPM) read count quantification from the binary alignment map (BAM) file was performed using HTSeq (v0.11.3), SAMtools (v1.10) and StringTie (version 2.1.1). Alignment, QC, and read-count quantification were performed using a nextflow (v#) pipeline hosted on a docker instance of Ubuntu 18.04.

### Partial Least-Squares (PLS) regression

Nutrient usage and production for partial least squares (PLS) regression was defined as the difference between the final and initial concentration of each quantified medium component in the rAAV production stage. Cell growth was defined as the difference between the peak cell density and the initial cell density in the production stage. Cell death was computed as the difference between the peak cell density and the final cell density. To minimize the skewing by larger magnitudes (e.g., glucose and lactate), the data were normalized by centering around the mean nutrient usage/consumption and dividing by the standard deviation. PLS analysis was performed in MATLAB^TM^ using the normalized nutrient data as the predictor and normalized rAAV titer was the response. Seven PLS components were used based on a 95% explained variance cutoff. The complete data required for performing the PLS regression is available in Supplementary Table ST1.

### Computing uptake and secretion rates

Uptake and secretion rates were computed using non-linear regression as previously described^8^. For the growth stage, the cells were assumed to be in a single metabolic state throughout the process. Therefore, a single set of extracellular fluxes were identified that minimizes the variance weighted squared deviation of predicted concentrations from measured data. Predicted concentrations were computed by solving the system of ODEs as described in Equations (1) and (2).

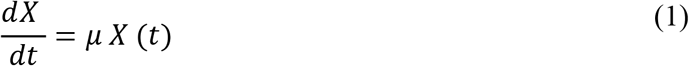

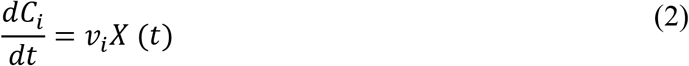

In the above equations, *X*(*t*) and *C_i_*(*t*) represent the cell density and concentration of metabolite *i* at time *t*, espectiielyy. *μ* and *v_i_* represent the growth rate and the uptake or secretion flux of metabolite *i*, respectively.

For the production stage, three metabolic states were assumed between the following time intervals: 0hpi – 6hpi, 6hpi – 24hpi, and 24hpi – 48hpi. Equations (1) and (2) were adjusted to account for time interval *k* using Equations (3) and (4).

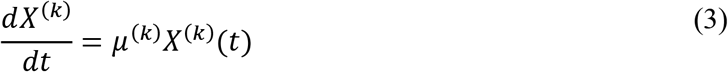

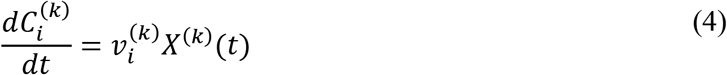

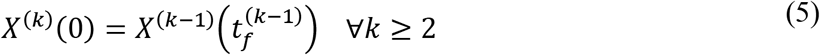

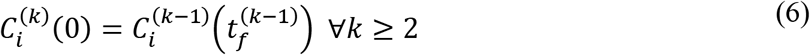

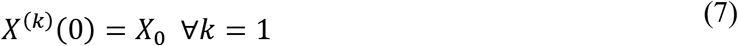

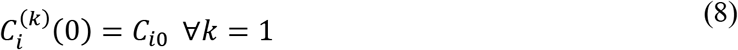

The above nonlinear least-squares problems were solved using the *fmincon* function within the Optimization Toolbox in MATLAB^TM^. Statistical acceptance was evaluated using the χ^2^-goodness of fit test with degrees of freedom defined as the difference between the number of data points and the number of parameters. All fluxes computed on a per cell basis were converted to a per gram dry weight (gdw) basis using a dry weight of 400 pg/cell. The uptake and secretion rates in mmol/gdw-day are provided in Supplementary Table ST2.

### Transcriptomic overlay onto metabolic models

Context-specific metabolic models for all clones and time points were constructed by overlaying gene expression data and extracellular fluxes onto the human metabolic reconstruction, Recon2.2^21^, as described earlier^22^. RNA-Seq data were normalized using the RLE normalization described in the DESeq2 protocol^44^ and log_10_ transformed. Next, ubiquity scores^23^ were computed using the gene-specific min-max-mean thresholding^24^ using only genes present in Recon2.2. Genes with a ubiquity score less than 1 were flagged as non-core reactions for removal and the remaining genes were flagged as core reactions for retention in context-specific models. Model pruning was performed using the modified mCADRE algorithm^23^ with an emphasis on ensuring compatibility with measured phase-specific uptake and secretion rates^22^. The *η* parameter was set to 0 to prohibit removal of reactions associated with highly expressed genes. Pairwise comparison of model content was quantified using the Jaccard index as defined in Equation (9).

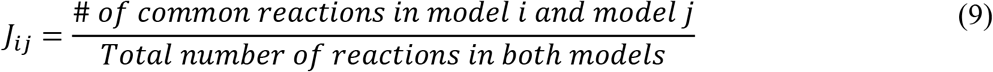

*J_ij_* assumes a value between 0 and 1 with 0 indicating the models are completely dissimilar to each other and a value of 1 indicating that the models are identical. For the reaction content of each context-specific model, see Supplementary Table ST3.

### Inferring pathway activities using CellFie

Pathway usage based on transcriptomic data was estimated using CellFie^24^. RLE-normalized gene expression data was thresholded using the min-max-mean approach. Task scores for 195 curated metabolic functions in Recon2.2 were computed using the thresholded gene expression data. For the complete metabolic task scores for each clone and time point, see Supplementary Table ST4. Variances in task scores were assessed using Principal Component Analysis (PCA) and Linear Discriminant Analysis (LDA) to elucidate the factors associated with master well differences and changes in rAAV productivity.

## Supplementary Figures

**Supplementary Figure S1:**
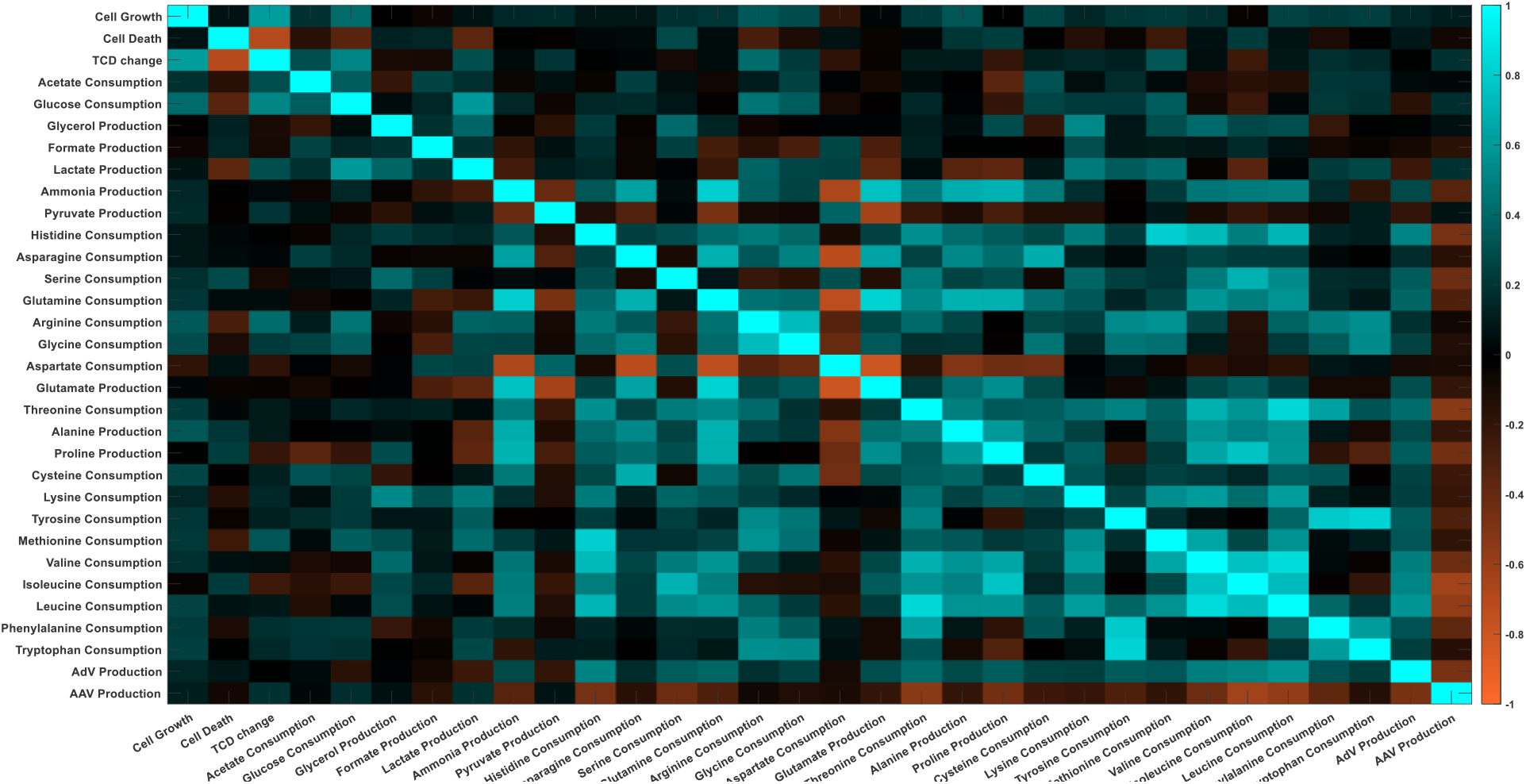
Heatmap of Spearman correlation coefficients showing the co-utilization and trade-off patterns between pairs of measured quantities.

**Supplementary Figure S2:**
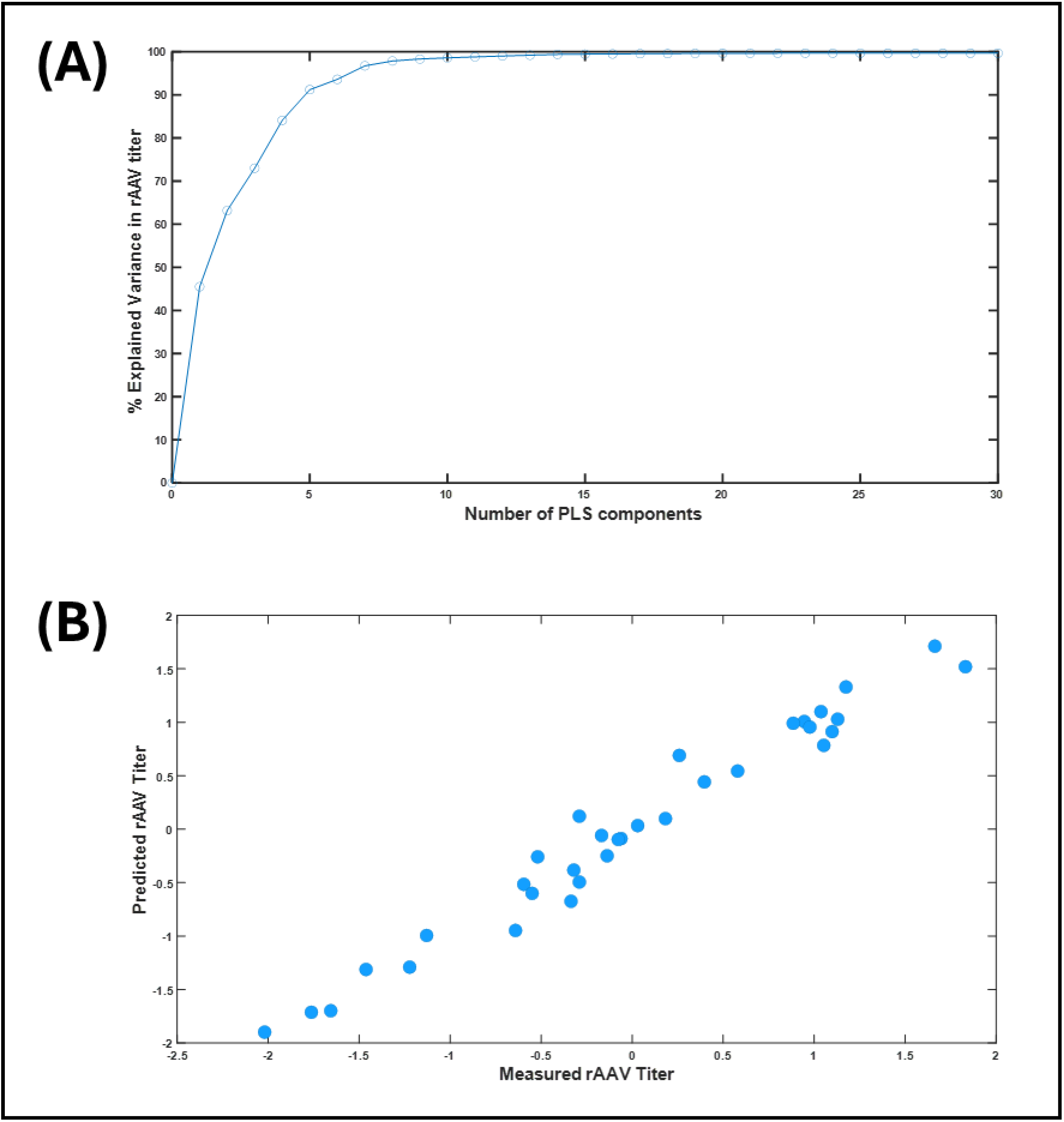
Supporting figures for PLS regression. (A) Explained variance as a function of number of components. (B) Agreement between predicted and observed rAAV titers.

**Supplementary Figure S3:**
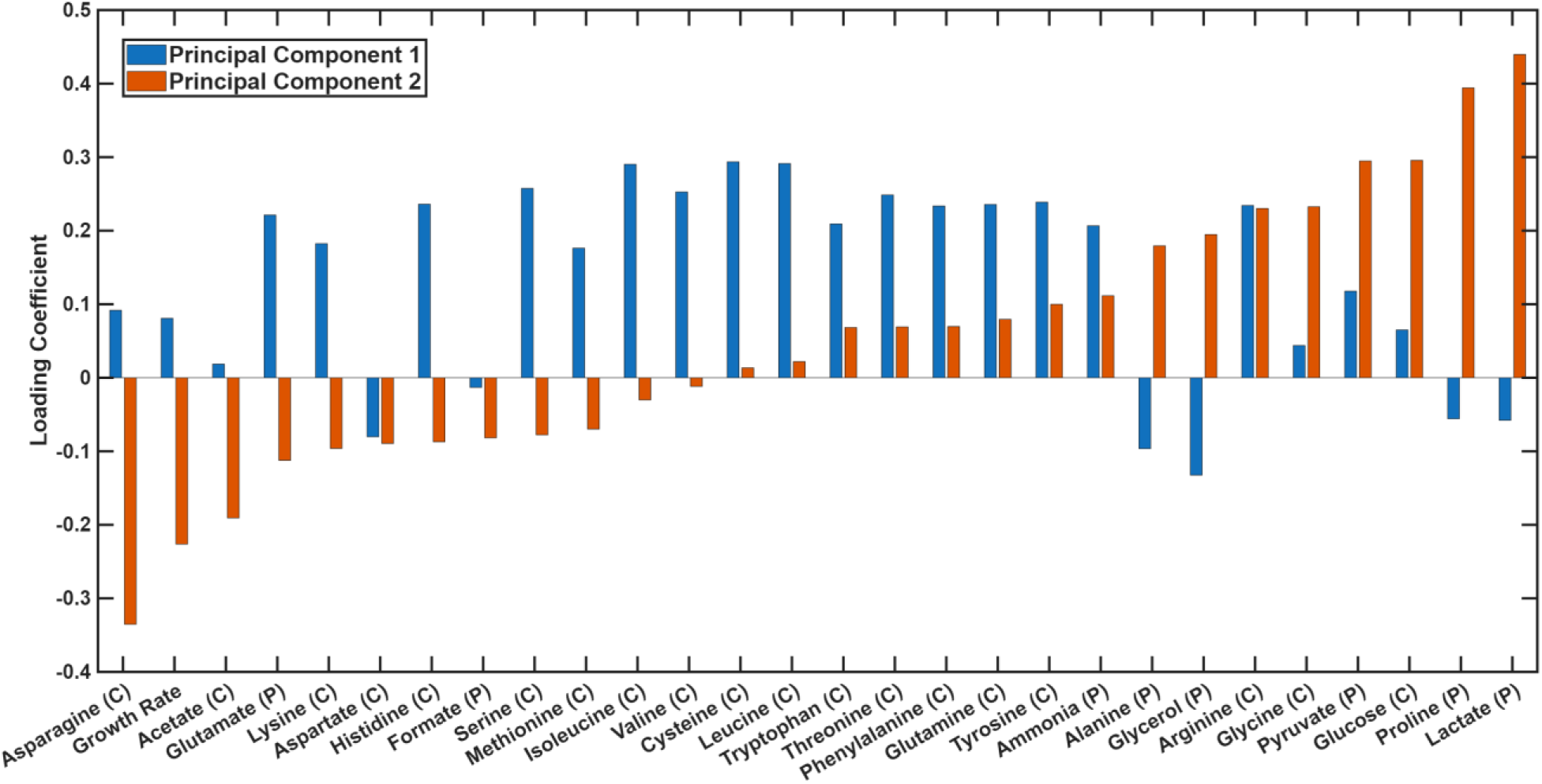
Loading Coefficients for the first two principal components for the PCA plot shown in Figure 3B.

